# Flow cytometry-based biomarker assay for *in vitro* identification of heat tolerance conferring coral symbionts

**DOI:** 10.1101/2022.11.21.517290

**Authors:** Patrick Buerger, Marcin Buler, Heng Lin Yeap, Owain R Edwards, Madeleine JH van Oppen, John G Oakeshott, Leon Court

## Abstract

Corals’ tolerance to high temperature stress largely depends on their symbiotic microalgae (Symbiodiniaceae). However, the contributing microalgal traits are largely unclear. Here we compare the *in vitro* cellular profiles of seven *Cladocopium C1^acro^* microalgal strains (derived from the same ancestral strain) during a four-week exposure to 27°C or 31°C. One was an unselected wild-type strain (WT), three were selected at 31°C for nine years and shown to confer thermal tolerance on the coral host (SS+) and three others were similarly selected but did not confer tolerance (SS-). Flow cytometry was used to measure the intracellular stress indicators reactive oxygen species (ROS), reduced glutathione (rGSH) and mitochondrial-membrane potential (ΔΨm), as well as cell size/shape and photosynthetic pigments. Cell densities and photosynthetic efficiency (ΦPSII, Fv/Fm) were also measured. WT showed the highest levels of intracellular ROS and ΔΨm, lowest rGSH and largest cell sizes at both temperatures. SS+ strains had the lowest ROS and highest rGSH values and a unique pattern of correlations among parameters at 31°C. Our results support previous reports implicating the role of microalgal ROS, ΔΨm and rGSH in holobiont thermal tolerance and suggest flow cytometry is a useful pre-screening tool for identifying microalgal strains with enhanced thermal tolerance.

## INTRODUCTION

Coral reefs are deteriorating rapidly across the world due to the increased incidence, severity and duration of marine heatwaves (Hughes *et al*., 2017). These extreme events disrupt the symbiosis between coral and endosymbiotic microalgae (Symbiodiniaceae) causing the loss of the algae from the coral tissue in a process known as coral bleaching (Glynn, 1996). Corals depend on photosynthates translocated from the Symbiodiniaceae for their nutrition, and without timely recolonisation of Symbiodiniaceae the coral is likely to die. Mass bleaching events, in which whole regions of reef bleach severely, have been observed increasingly frequently in many parts of the world over recent years due to summer heatwaves (Hoegh-Guldberg, 1999; Hughes *et al*., 2018). For example, on the Great Barrier Reef (GBR) mass bleaching events have been recorded frequently in recent years, including 2016, 2017, 2020 and 2022, whereas fewer of such events had been recorded there previously (e.g., in 1998, 2002 and 2006). The impacts of the recent events have increased in severity and footprint (Hughes *et al*., 2021), reducing coral recruitment and shifting community diversities, which is predicted to also impact ecosystem functioning (Cheung *et al*., 2021). Targeted intervention strategies could support coral adaptation to climate change if carbon emissions are simultaneously reduced (Anthony *et al*., 2017; Bay *et al*., 2019) but many of these strategies depend on understanding the molecular mechanisms underpinning thermal tolerance (van Oppen *et al*., 2017).

One of the main drivers of coral bleaching is assumed to be excess production of reactive oxygen species (ROS) by the Symbiodiniaceae during heat stress (Suggett *et al*., 2017). Excess ROS can overwhelm the antioxidant capacities of both the symbiont and the coral, leading to cellular malfunctions and ultimately coral bleaching (Downs *et al*., 2002; Weis, 2008; Nielsen *et al*., 2018). Accordingly, declines in the functionalities of both microalgal and host organelles such as chloroplasts and mitochondria can be early indicators of heat stress (Roberty and Plumier, 2022). For example, the breakdown of chloroplast functionalities can be measured as declines in chlorophyll content and the photosynthetic efficiency of photosystem II (as measured by maximum and effective quantum yields), while changes in mitochondrial membrane potential (ΔΨm) can be potential stress indicators (Dourmap *et al*., 2020; van Aken, 2021). Potentially counteracting these effects, however, antioxidant defence mechanisms such as superoxide dismutase enzymes and the reduced form of glutathione (rGSH) can help maintain an equilibrium between ROS production and scavenging, thus minimising cellular damage and sustaining the natural molecular signalling function of ROS (Hasanuzzaman *et al*., 2020). Notably, differences between Symbiodiniaceae species and strains have been found in the activities of several of these elements of ROS production, damage and scavenging, and these differences could be important sources of variation in the tolerance of these symbionts to elevated temperatures *in vitro* and *in hospite* (Suggett *et al*., 2017).

One promising study system for examining the physiological basis of variation in the heat tolerance properties of the Symbiodiniaceae involves a set of closely related strains of *Cladocopium* C1^acro^ developed through experimental evolution. Heat-evolved *Cladocopium* C1^acro^ strains, referred to as heat-evolved strains (SS strains) and their wild-type counterparts (WT strains) were established from the same mother culture in 2011 and kept at 31°C and 27°C respectively since their establishment (Chakravarti *et al*., 2017). The ten SS strains show higher tolerance to 31°C, as reflected in higher growth rates, maintenance of photosynthetic efficiency and lower net ROS compared to the WT strains (Buerger *et al*., 2022). Some of the SS strains (hereafter SS+ strains) also enhanced thermal bleaching tolerance of coral larvae and juveniles, although other SS strains (hereafter SS-) did not. Specifically, larvae of the coral *Acropora tenuis* in symbiosis with one of the SS+ strains (SS1, SS7 or SS8) were found to maintain symbiont numbers in their tissues during a short term heat stress experiment while larvae with SS- or WT strains bleached (Buerger *et al*., 2020) and juveniles of the coral *A. tenuis* in symbiosis with a SS+ strain (SS1) showed higher thermal tolerance than juveniles with a WT strain (Quigley and van Oppen, 2022).

Flow cytometry has been used in coral related research mainly to enumerate Symbiodiniaceae cells in samples (e.g., Lee et al., 2012; Krediet et al., 2015; McIlroy et al., 2016). Only very few studies have used flow cytometry to assess Symbiodiniaceae cells in response to heat stress (Lee et al., 2012; Delamare-deboutteville et al., 2020; Maruyama et al., 2022). However, to the best of our knowledge, a flow cytometry assay has not been applied to identify Symbiodiniaceae strains with different thermal tolerance physiologies or to find biomarkers for enhanced thermal tolerance of coral associated microalgae.

Hence, the objective of the current experiment was to identify traits that distinguish strains with different thermal tolerances and capacities to confer their tolerance to the coral host (comparing three SS+ strains to the SS- and WT strains). To this end, the three SS+ strains plus three SS- and one WT strain were characterised *in vitro* for nine physiological and morphological traits previously implicated in thermal tolerance over a 28-day period at ambient (27°C) and elevated temperatures (31°C). The parameters assessed were: cell densities (CellDen) through automated cell counts; photosynthetic efficiency with maximum quantum and effective quantum yields (ΦPSII, Fv/Fm) through pulse-amplitude modulation chlorophyll fluorometry; and flow cytometry-based parameters such as intracellular reactive oxygen species (ROS), mitochondria potential (ΔΨm), reduced glutathione (rGSH), photosynthetic pigments (mainly chlorophyll; Chl, see Experimental Procedures), plus cellular size and complexity as measured by forward and side scatter (FSC and SSC, respectively; (Shapiro, 2005), Figure S1).

## Material and Methods

### Microalgae strains

Seven *Cladocopium* C1^acro^ strains were obtained from the Australian Institute of Marine Science in May 2021 (culture collection numbers: SCF-055.10 for the wild-type strain WT10, SCF-055.01, SCF-055.07, and SCF-055.08 for the SS+ strains SS1, SS7 and SS8 respectively, and SCF-055.03, and SCF-055.05, SCF-055.09 for the SS-strains SS3, SS5 and SS9, respectively). The strains were established in an assisted evolution experiment that started in 2011, in which the temperature was increased for the SS strains from ambient 27°C to elevated 31°C over two months in a step-wise fashion and the best growing strains were transferred to the next step. Then, the strains were cultivated at 31°C on a permanent basis and showed enhanced *in vitro* thermal tolerance in several performance assessments (Chakravarti *et al*., 2017; Buerger *et al*., 2022). Some of the strains, here termed SS+, also conferred their thermal tolerance *in hospite* when in symbiosis with coral larvae (Buerger *et al*., 2020) and juvenile corals (Quigley and van Oppen, 2022). In comparison, the wild-type strain has been maintained at 27°C since 2011.

### Experimental set up

Throughout the current experiment (conducted in 2020), the strains were incubated in temperature-controlled rooms at either ambient (27°C) or elevated temperatures (31°C) with 50% relative humidity, using LED lights at a light intensity of 31 to 33.5 PAR on a 12:12 light:dark cycle. Each culture was provided 0.2 μm of filtered IMK medium prepared from Mili-Q water and Red Sea Salt (Red Sea/cat: 398-01333) and enriched with Daigo’s IMK Medium for Marine Microalgae (cat: 398-01333, Nihon Pharmaceutical Co).

To obtain enough biological material for the experiment, the seven strains were each cultured for 3 months at 27°C in 100 mL media (175 cm^2^ growth flasks, cat: NUN159910, Thermo Fisher Scientific) and then split into 25 cm^2^ flasks for the assays, seeded at 3 * 10^5^ cells / mL. At each time point, a total of five replicate assay flasks were sampled from the six SS strains and six replicate assay flasks from the WT10 strain (total number of replicates per sampling time point = 36). To minimise evaporation, the assay flasks were arranged and stored in clear plastic boxes, each with a combination of five SS strain assay flasks and the WT10 control assay flask. The boxes with the assay flasks were placed at 27°C to allow them to acclimate for two days after subculturing. To minimize variation in light exposure within the respective temperature-controlled rooms, all box positions were rotated on the shelf twice per week and assay flask positions were randomized within a box at each subsampling time-point.

The nine response variables were measured on subsamples of each flask at Days 4, 11, 18 and 28. Each time, the microalgal cells were detached from the assay flask surface using a sterile disposable cell scraper (cat: 83.3951, Sarstedt) and resuspended uniformly by gentle pipetting using an S1 Pipet Filler (cat: 9531, Thermo Fisher Scientific) fitted with a sterile 0.45 μm hydrophobic filter (cat: MAT9057, Thermo Fisher Scientific) and a sterile disposable 10 mL serological pipette (cat: 357551, Corning). A total of 6.5 mL was removed per assay flask at each sampling time and transferred to a sterile 15 mL tube (cat: 430766, Corning) until split proportionally for each of the following measurements.

### Cell densities

Cell density readings were obtained on a TALI image-based cytometer (cat: T10796, Thermo Fisher Scientific). Two technical replicate readings of 25 μL were made per assay flask and averaged to give a final estimate. TALI cytometer settings for each reading were: viability assay, 20 fields, cell gating 5 to 14 μm, sensitivity setting 6; circularity setting 8. The cell count measurements were then used to normalize each strain’s subsamples to 0.4 * 10^6^ cells / mL prior to the staining and hence also for the following flow cytometry measurements.

### Cell staining

After density normalization, the cells were transferred to 2 mL tubes (cat: 0030120094, Eppendorf) in 1.5 mL aliquots for Unstained Control cells, CellROX Orange staining and MitoTracker Orange staining workflows and in 1 mL aliquots for the Vita-Bright-48 staining workflow. Cells in the former 1.5 mL aliquots were pelleted at 2,600 g for 5 minutes and washed in 1.5 mL of fresh media, pelleted again as before and concentrated slightly by resuspending in 1.2 mL of fresh media to enrich for microalgal cells and optimise staining conditions. The 1.2 mL cell suspensions were then split into three separate 2 mL tubes, with one aliquot of 0.2 mL for unstained control cells, one of 0.5 mL for CellROX Orange staining and one of 0.5 mL for MitoTracker Orange staining (Table S1). The cells were then subjected to the respective staining protocols below and measured for their mean fluorescence intensity (MFI) on the Attune NxT Flow Cytometer. Cells in the latter 1 mL aliquots were processed as described below in the Vita-Bright-48 workflow.

### Intracellular ROS staining with CellROX Orange

To measure ROS, 0.5 mL subsamples of cell culture normalized as above were resuspended by vortexing and then stained with CellROX Orange reagent (cat: C10443, Thermo Fisher Scientific). To prepare a 250 μM working stock of the staining solution, a 16 μl subsample of CellROX Orange reagent at 2.5 mM was diluted 1:10 by the addition of 144 μl of fresh media (as per manufacturer’s instructions). Staining of cells was synchronized across all tubes by pipetting 6 μl of the working stock into the lid of each 2 mL tube and then mixing all tubes in a tube rack at once by repeated inversion. The cells were then incubated for 60 minutes at room temperature in a dark cupboard (final stain concentration of 3 μM).

### rGSH staining with Vita-Bright-48

To measure intracellular reduced thiols, 1 mL subsamples of cell culture normalized as above were resuspended by vortexing and stained with Vita-Bright-48 contained in Solution 6 (cat: 910-3006, ChemoMetec; 1.34 mM for Vita-Bright-48; 0.75 mM PI) (Skindersoe *et al*., 2012). Staining was synchronized across all 2 mL tubes by first pipetting 10 μl of Solution 6 into the lid of each 2 mL tube and then mixing all tubes in a tube rack at once by repeated inversion. The cells were then incubated for 90 minutes at room temperature in a dark cupboard (final stain concentration of 13.4 μm for Vita-Bright-48 and 7.5 μm for PI). After staining, the microalgae were washed twice by pelleting the cells each time at 2,600 g for five minutes and then resuspending in 1 mL fresh media by briefly vortexing. The cells were pelleted one final time as above and then resuspended in 450 μl fresh media.

### ΔΨm staining with MitoTracker Orange

Mitochondrial membrane potential ΔΨm) was assessed using the fluorescence-based stain MitoTracker Orange within live microalgal cells. 0.5 mL subsamples of cell cultures normalized as above were dispensed into 2 mL tubes (cat: 0030120094, Eppendorf) and stained with MitoTracker Orange CMTMRos (cat: M7510, Thermo Fisher Scientific). Each tube of CMTMRos stain contained 50 μg lyophilized solid, which was prepared for use by reconstituting to 200 μM with the addition of 585 μl DMSO. A 5 μL subsample was then diluted 1:40 by the addition of 195 μL DMSO to produce a 5 μM working stock. Staining of microalgal cells was synchronized across all tubes by pipetting 5 μl of 5 μM CMTMRos into the lid of each 2mL tube and then mixing all tubes in a tube rack at once by repeated inversion. The microalgal cells were then incubated for 80 minutes at room temperature in a dark cupboard (final stain concentration of 5 nM, Table S1).

### Flow Cytometry

Microalgal cells were acquired with the Attune NxT Flow Cytometer and analysed with (Attune NxT software, Thermo Fisher Scientific). All staining protocols were optimized to ensure stable signal retention during sample preparation and acquisition. Only single, photosynthetic pigments-positive cells were selected with specific gating configurations (Figure S1). Details of instrument settings, number of cells measured, acquisition and drawing volumes and peak emissions/bandwidths for the filters used and the excitation/emission maxima of the three stains are given in Table S2. The median fluorescence intensity (MFI) of pigments-positive cells and their side and forward scatter heights, SSC-H and FSC-H, respectively, were determined in unstained controls (UC), while samples stained with the respective dyes were treated independently and acquired separately to avoid spectral crossover. Preliminary analyses showed no relationship between cell size and stain intensity for any of the stains.

Due to a relatively high background signal and/or weak dye signal in samples treated with Vita-Bright-48 and CellROX Orange dyes respectively, values for these stains were calculated by subtracting the measured unstained controls (MFI_UC_) from the stain’s MFIs (Figure S1). In contrast, a distinctive peak separation in MitoTracker Orange CMTMRos stained samples allowed us to measure mitochondrial potential in cells without MFI*_UC_* subtraction.

The Attune was configured with three lasers, one of excitation 405 nm for measuring rGSH and Chl, one of excitation 488 nm for measuring FSC and SSC, and one of excitation 561 nm for measuring ROS and ΔΨm. In order to ensure consistent measurements between samples and to avoid potential instrument blockages, we performed a rinse step after every 6 samples. Note that we do not know which specific photosynthetic pigments contributed to our Chl measurements because chlorophyll a, b and c and carotenoids have overlapping absorption spectra and are all excited by the 405 nm laser.

### Photosynthetic efficiency

At each time point, photosynthetic efficiency was assessed using an imaging pulse-amplitude modulation chlorophyll fluorometer (IPAM M-series, Walz, Germany). Measurements of maximum quantum yield [Y(II) = Fv/Fm] (Murchie and Lawson, 2013) were performed after midday on samples that were still dark-adapted from the night cycle. Effective quantum yield measurements [ΦPSII = ΔF/Fm’] (Murchie and Lawson, 2013) were taken after a five-minute acitinic light exposure in order to light-adapt the samples to assess the photochemical efficiency of photosystem II. For both variables, measurements at three circular locations were averaged as technical replicates for each flask. The following instrument setting were used: measurement intensity, 6; saturation pulse, 10; gain, 2; damping, 2. The data for each strain with means across replicates are given in Figure S2, while the raw data is available in the supplementary data.

### Statistical analysis

All nine response variables were normalised to unit ranges. We then ran PERMANOVA (R version 4.1.1, package *vegan*) for each individual variable against a full interaction model for the factors temperature, sampling time and strain group. MANOVA and ANOVA were not used because the data for all variables failed normality, heteroscedasticity and outlier tests (from the R packages *mvnormtest, biotools* and *mvoutlier* respectively). Pearson correlations were calculated between the respective parameters and visualized as a matrix. We then performed linear discriminant analyses (LDAs) on the normalised variables to visualise the differences between strain groups across temperatures and sampling times. Pairwise plots to demonstrate group separations considering minimal numbers of parameters were also generated.

## RESULTS

### Changes in individual variables

None of the three non-flow cytometry measurements CellDen, ΦPSII and Fv/Fm differentiated the three types of strains at 27°C, but all three differentiated them at 31°C from Day 4 (Figures 1A, S2). At 27°C, CellDen increased to a plateau by ~11 Days while ΦPSII and Fv/Fm values decreased, particularly after ~11 Days. At 31°C, the CellDen of all three strain types only rose slightly up to ~11 Days and then fell, but the trajectory for SS+ was higher than that of SS-, which in turn was higher than that of WT. ΦPSII and Fv/Fm at 31°C again decreased in all three strain types but the decrease was less for the SS+ than the SS- or WT strains.

**Figure 1.**
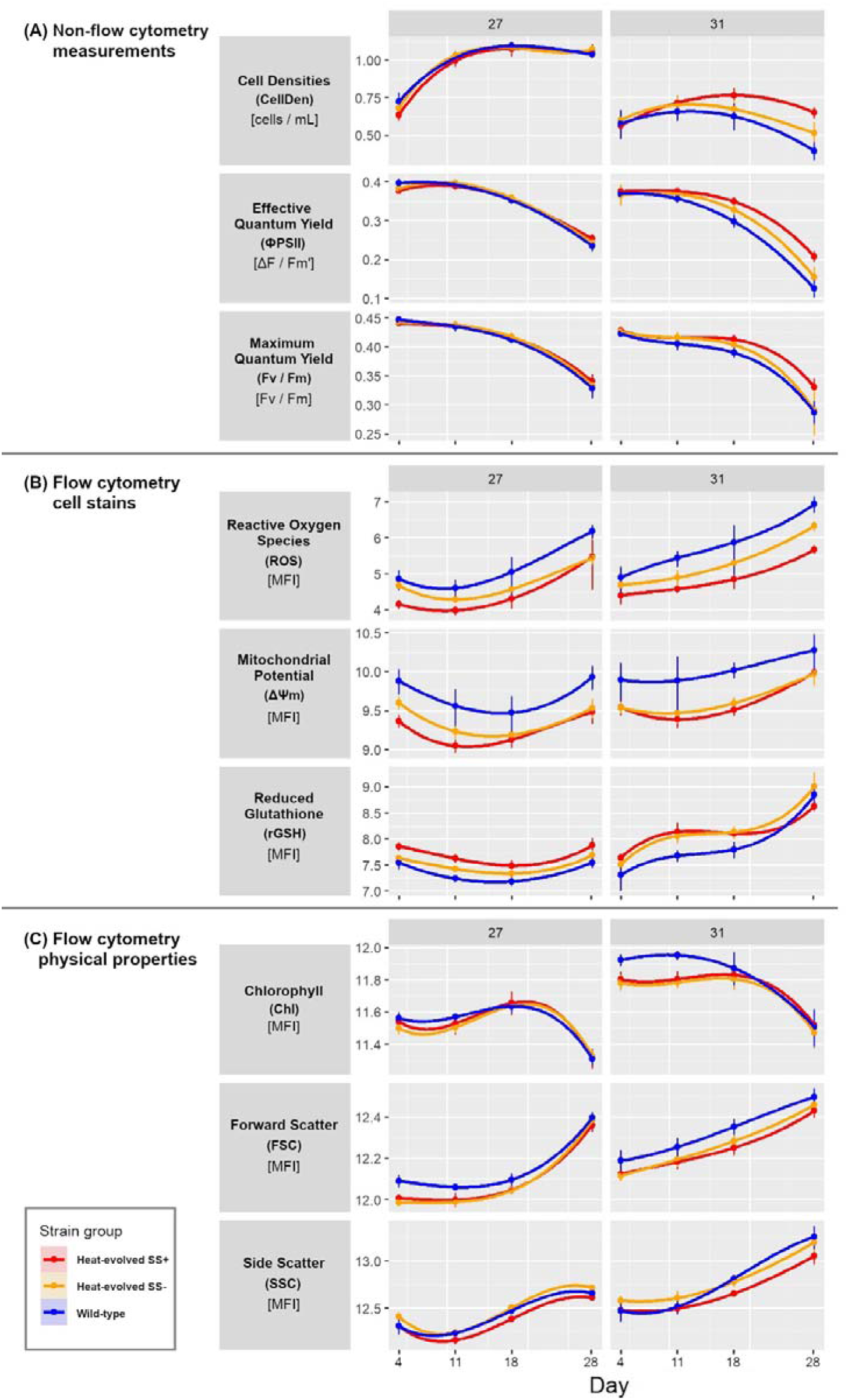
Progress curves for the three strain types for each variable and temperature. **(A)** Non-flow cytometer measurements of cell density, effective quantum yield and maximum quantum yield. **(B)** Flow cytometry cell dye measurements of reactive oxygen species (CellROX Orange), mitochondrial potential (MitoTracker Orange CMTMRos) and reduced glutathione (Vita-Bright-48), all displayed as mean fluorescent intensity [MFI]. **(C)** Flow cytometry measurement of the physical properties photosynthetic pigments, and cell size and shape (forward and side scatter respectively). All variables are plotted on natural log scales. Error bars show 95% confidence intervals based on 5 or 6 replicates and assuming t-distributions. Measurements were taken on Days 4, 11, 18 and 28.

In contrast, the three dye-based flow cytometry measurements ROS, ΔΨm and rGSH revealed differences between the three types of strains at both temperatures (Figure 1B). At 27°C the values for all the measures dropped slightly initially and then rose again, with SS+ always lower than WT for ROS and ΔΨm and higher for rGSH. SS- was generally intermediate for all three variables but moved closer to SS+ for ROS and ΔΨm over time. At 31°C the values for all three variables showed a stronger upward trend over time. SS+ values were again consistently lower than WT values for ROS and ΔΨm. rGSH values were also again higher initially for SS+ than WT but in this case the difference disappeared over time. SS- was again consistently intermediate for ROS but similar to SS+ for the other two variables.

Relatively few differences between the strain types were evident in the flow cytometry measurements of physical cell properties (FSC, SSC) and photosynthetic pigments (Chl) at either temperature (Figure 1C). There was little change in any of the three measures over the first three time points at 27°C, but by Day 28 Chl had dropped and FSC had risen considerably, and SSC had risen to a lesser extent. While SS+ and SS- were initially lower than WT for FSC, that difference had largely disappeared by Day 28. The directions of the changes over the course of the experiment for each measure at 31°C were similar to those at 27°C but the relationships between the strains changed over time for both Chl and SSC. Chl values were initially lower for SS+ and SS- than WT but the difference disappeared by Day 14. SS- values were always higher than SS+ values for SSC but WT values were initially similar to SS+ but similar to SS- at Days 11 and 28. On the other hand, SS+ and SS- showed consistently lower FSC values than WT at 31°C throughout the experiment.

In sum, all nine measures changed during the experiment, with the trajectories of several differing between temperatures and most differing between strain types, particularly between SS+ and WT, for at least one of the temperatures. Of the key parameters proposed from earlier work (see Introduction), ROS and mitochondrial potential showed relatively minor differences between temperatures but relatively large differences between strain types, while rGSH showed greater differences between temperatures but less consistent differences between strains. Differences also occurred in photosynthetic pigment amounts and photosynthetic efficiencies, although the patterns of change differed; there were clear drops in all three measures in the last time interval, particularly at 31°C, as well as strain type differences at that interval and temperature for photosynthetic efficiency but not photosynthetic pigment content. Cell density increased early and cell shape changed late in all three strain types at 27°C, whereas at 31°C cell density did not show marked temporal change, but clear strain type differences emerged in all three measures, particularly cell density.

Notably also, the changes in the last time interval for many of the variables at both temperatures suggested the cells were becoming stressed by Day 28 which could be related to the age of the cultures and likely nutrient depletion in the media.

### Permutational analysis of variance

As expected from the progress curves, three-way factorial permutational analyses of variance (PERMANOVAs) showed large main effects of time and temperature and smaller but still significant effects of strain type on all nine parameters (Table 1). However, at least two of the four interaction terms were also significant for all the variables, and in all cases at least one of them involved strain type. The combined contributions of main or interaction terms involving strain type to the total variance were ~28% for ROS and ~23% for ΔΨm but less than 10% for all the other variables. For most of the variables, including ROS, the main effect of strain type contributed less than half of their respective combined contributions but for ΔΨm it accounted for most of it (~21 of the 23%).

**Table 1.** Analyses of variance. Sums of squares are expressed in percentages [%s] for each analysis. Significant terms for each variable are highlighted in grey. * p < 0.05; ** p = 0.01; *** p < 0.001. Df = degrees of freedom.

| Term | Df | CellID | Fv/Fm | ΦPSII | ROS | $\Delta\Psi_m$ | rGSH | Chl | FSC | SSC |
| --- | --- | --- | --- | --- | --- | --- | --- | --- | --- | --- |
| Time | 3 | 21.6*** | 78.1*** | 82.8*** | 41.3*** | 24.9*** | 25*** | 36.3*** | 68.9*** | 49.2*** |
| Temperature | 1 | 53.8*** | 3.7*** | 4.5*** | 7.3*** | 17.7*** | 19.4*** | 36.2*** | 19*** | 26.6*** |
| Strain | 2 | 0.3* | 0.7** | 0.8*** | 12.3*** | 20.8*** | 1* | 1.7*** | 2*** | 2.7*** |
| Time : Strain | 6 | 1*** | 0.9* | 1*** | 9.4*** | 0.3 | 1.9* | 0.9 | 0.2 | 0.8** |
| Time : Temperature | 3 | 12.9*** | 0.4 | 1.9*** | 5.2*** | 6.8*** | 19.5*** | 3*** | 2.3*** | 8.8*** |
| Strain : Temperature | 2 | 0.9*** | 0.5* | 1*** | 2.6*** | 0.6 | 1.1** | 0.8* | 0.1 | 0.3* |
| Time : Temp : Strain | 6 | 0.2 | 0.3 | 0.1 | 3.5*** | 0.9 | 2.1** | 0.3 | 0.1 | 0.5 |
| **Residuals** | **264** | 9.4 | 15.6 | 7.9 | 18.4 | 28 | 30 | 20.8 | 7.3 | 11.1 |

In combination with the progress curves, these analyses suggest that the major differentiators among strain types were differences in ROS and ΔΨm, with expression of the former also heavily dependent on time and temperature but the latter less dependent on those other factors. However, the analyses also confirm that the other seven parameters also show some effect of strain type, much of it temperature- and/or time-dependent.

### Correlations among measurements

To better understand the relationships among the variables, pairwise correlational analyses (Pearson’s r) on various sets of the data were carried out. The first of these analyses included all the data (Figure 2A). The strongest positive correlations overall were between Fv/Fm and ΦPSII and between FSC and SSC (r = 0.92 and 0.85 respectively), which was expected given that the first two of these both measure photosynthetic efficiency and the second two both relate to cell size/shape. Fv/Fm and ΦPSII were also both strongly negatively correlated with both FSC and SSC (r between −0.83 and −0.75), suggesting lower photosynthetic efficiency in larger cells. Otherwise, the largest correlations were positive ones between ROS and FSC and SSC (r = 0.82 and 0.78 respectively) and negative ones between ROS and Fv/Fm and ΦPSII (r = −0.70 and −0.69 respectively), suggesting that the larger cells with compromised photosynthetic efficiency also had greater ROS.

**Figure 2.**
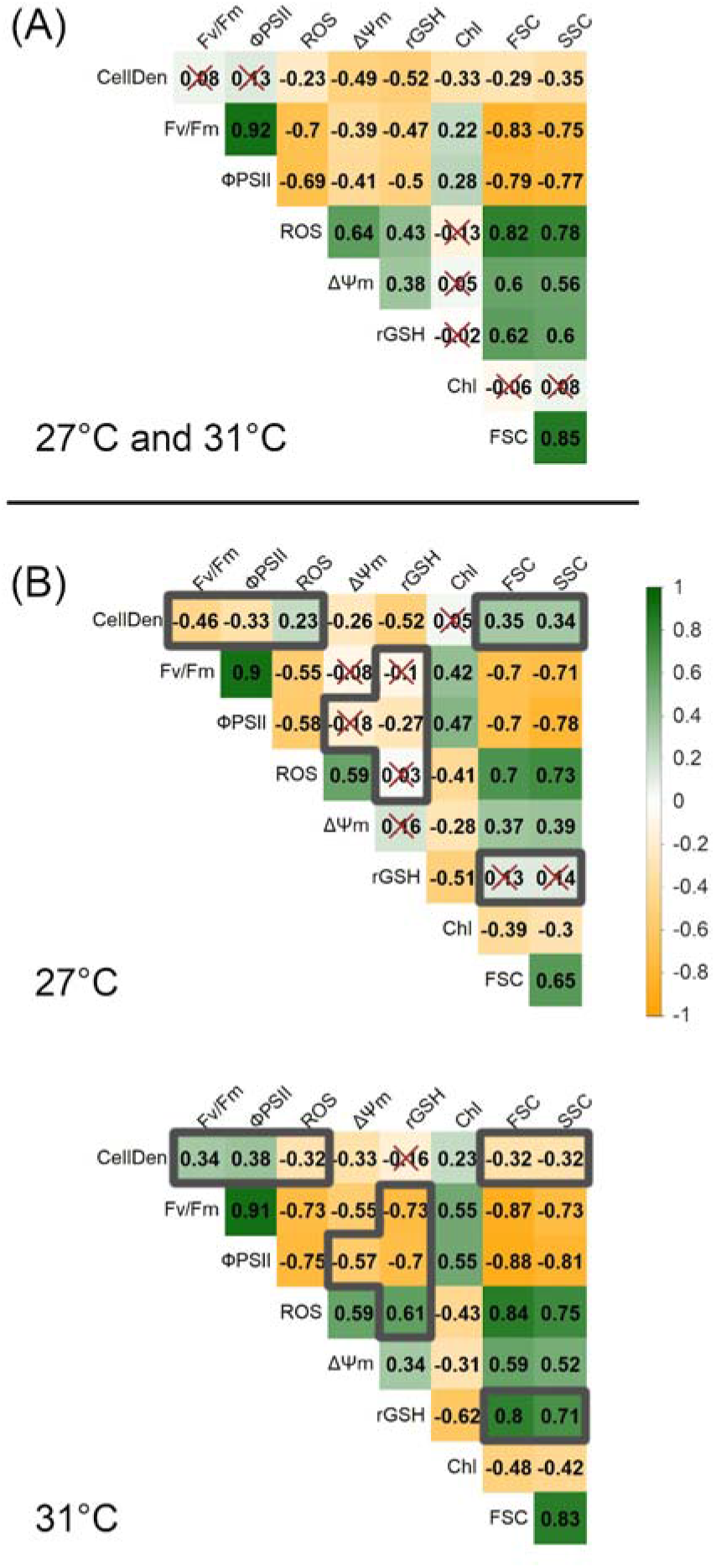
Pairwise Pearson correlation coefficients (r) between parameters. **(A)** shows the results for all data, combining 27°C and 31°C. **(B)** shows the results for each temperature separately. Grey borders mark r values that differ between 27°C and 31°C by ≥0.5. Crosses in both panels indicate r values < 0.2.

While several other positive and negative correlations were statistically significant, they generally explained less than half the variation in the constituent variables (i.e., r^2^ < 0.50), indicating that they provided considerable independent information.

Calculating the correlations separately for the two temperatures (Figure 2B) showed most of the correlations (31 out of 36) were larger in magnitude at 31°C. This suggests closer links between the differences in most of the variables at the elevated temperature. The only major exception was the correlation between CellDen and rGSH (−0.52 at 27°C *cf*. - 0.16 at 31°C). Considering also the magnitude, eleven of the correlations differed by more than 0.5 between the two data sets. In particular, at 31°C ROS was more strongly negatively correlated with Fv/Fm, ΦPSII and cell densities than at 27°C (r = −0.73, −0.75 and −0.32, cf −0.55, −0.58 and 0.32, respectively) and more positively correlated with rGSH (r = 0.61 cf 0.03, respectively), while rGSH was itself more negatively associated with Fv/Fm and ΦPSII (r = −0.73 and −0.70 cf −0.10 and −0.27 respectively). These differences are consistent with more severe effects of ROS at 31°C on photosynthetic output and cell health (as represented by cell densities) and an adaptive response in rGSH at that temperature to sequester ROS.

There were also much stronger positive associations of rGSH with FSC and SSC at 31°C than 27°C (r = 0.80 and 0.71 cf 0.13 and 0.14 respectively), suggesting that the larger cells at 31°C had higher content of rGSH than those at 27°C.

Other large differences between the 31°C and 27°C correlations included: reversals in the directions of the associations of CellDen with FSC and SSC (r = −0.32 and −0.32 cf 0.35 and 0.34 respectively) and CellDen with Fv/Fm and ΦPSII (r = 0.34 and 0.38 cf −0.46 and - 0.33 respectively), and stronger negative associations of ΔΨm with Fv/Fm and ΦPSII (r = - 0.55 and −0.57 cf −0.08 and −0.18). These differences suggest fundamental changes in the relationships between cell densities, cell size, photosynthetic efficiency and mitochondrial potential during the experiments at the two temperatures.

Finally, we computed the correlations among the measured parameters separately for both the two temperatures and the three strain types. The correlation matrices for the WT, SS- and SS+ strains at 27°C were very similar overall (Figure 3A). The largest differences were found in the correlations between CellDen and FSC (WT: 0.11, SS-: 0.54, and SS+: 0.25) and between ΔΨm and FSC (WT: 0.60, SS-: 0.11, and SS+: 0.34). Thus, at ambient temperature there was relatively little differentiation between the three strain groups in the relationships between parameters. However, several large differences were evident between the matrices for the three types of strains at 31°C (Figure 3B). Most notably, for the WT and SS- strains, CellDen were involved in several significant positive and negative correlations with other variables, whereas, for the SS+ strains, CellDen were not correlated with any other variable at this temperature. Together with the progress curves for the different variables (Figure 1), these results show that the SS+ strains maintained relatively stable cell densities at 31°C over the course of the experiment, despite significant changes in other variables. In contrast, CellDen (and some other variables) in the WT and SS- strain groups at 31°C changed much more over time.

**Figure 3.**
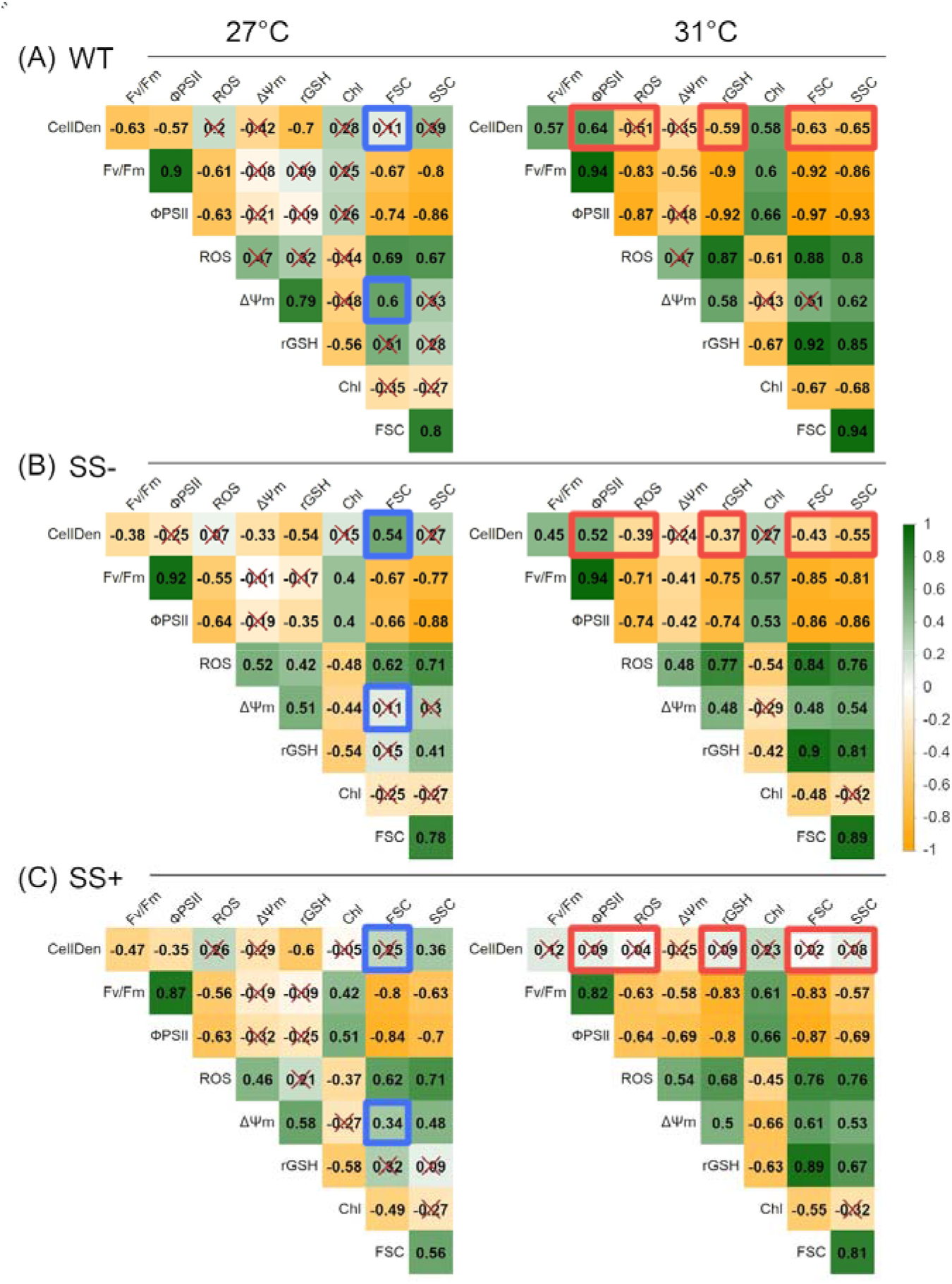
Pairwise Pearson correlation coefficients (r) between parameters for each strain type and temperature. **(A)** WT, **(B)** SS+ and **(C)** SS-. Blue and orange borders indicate r values that differ between strain types by ≥0.5 at 27°C and 31°C respectively. Crosses indicate r values < 0.2.

### Multivariate analyses

In view of the complex interactions between many of the parameters that were exposed in the previous analyses, we then used multivariate methods to assess their cumulative effects. We dropped FSC and ΦPSII at the outset of these analyses because they had shown broadly similar behaviours to SSC and Fv/Fm respectively in the previous analyses. A principal component analysis (PCA) of the seven retained variables found that most (94.6%) of their variation was accounted for by five principal components (Table S3). These five components were then used as the input for a linear discriminant analysis (LDA).

As might be expected from the analyses above, LD1 and LD2 (explaining 66.1% and 23.3% of the between group variance respectively) distinguished the data for different times and temperatures respectively (Figure 4, Table S4). However, the three strain types were also separated, with all three distinguished at 27°C and SS+ distinguished from the other two at 31°C. In particular, the SS+ strains also changed less over time (i.e., showed a shorter Day 4 to Day 28 distance) at the higher temperature than did the other strains. There was also less variation among the three constituent SS+ strains (i.e., less dispersion around the centroids) than among the three SS- strains (Figure 4 and see also Figure S3).

**Figure 4.**
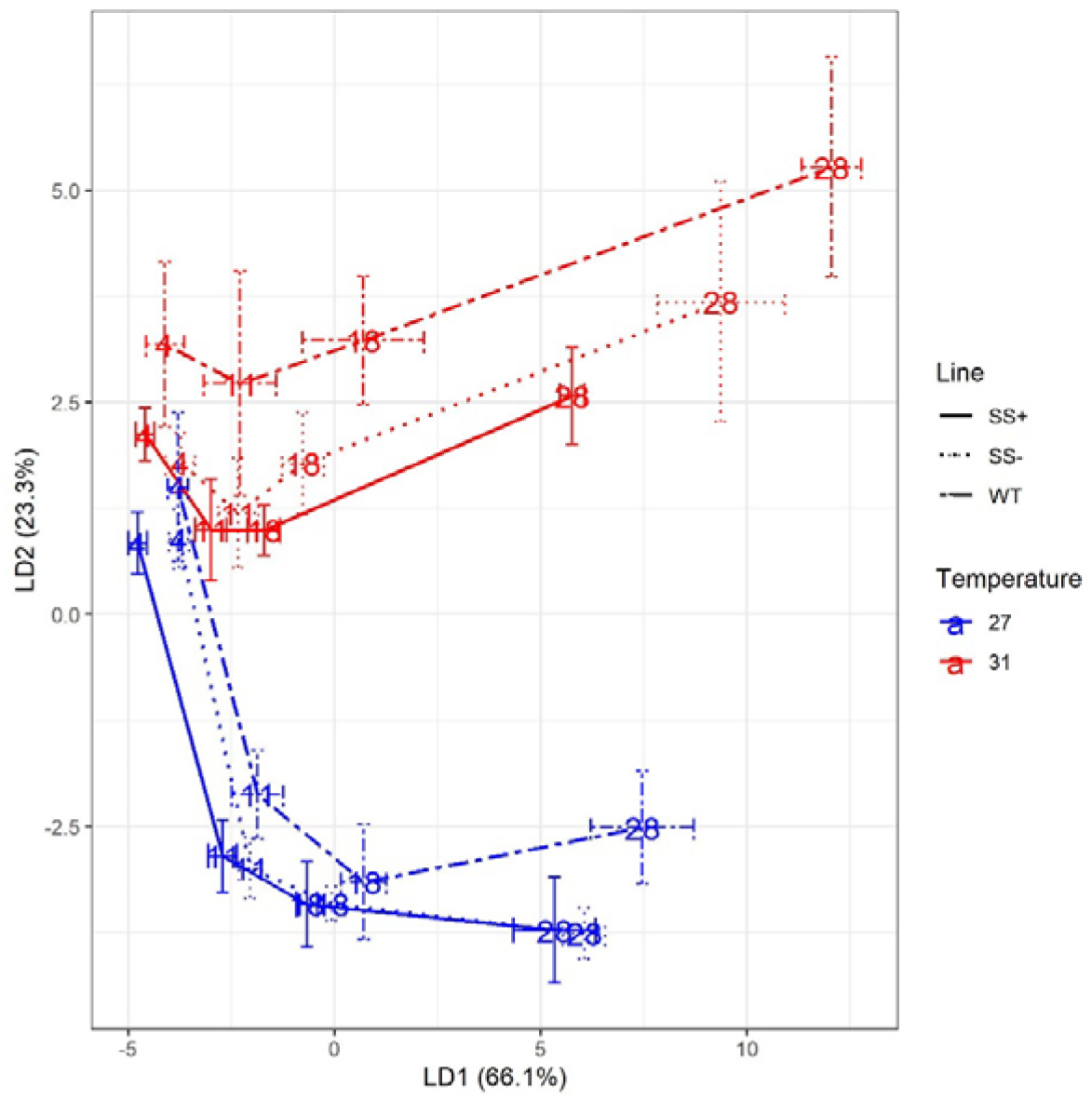
Plot of the two major linear discriminant functions for each strain type and temperature. The plot is based on all seven variables retained in the multivariate analyses. Mean values with their 95% confidence limits are given for each time point.

We next carried out separate LDAs on the data for each Day and with two subsets of the seven variables. Our goal here was to compare the resolving abilities of the five flow cytometer and two other measurements retained. We found that LDAs using the five flow cytometer measurements achieved similar levels of resolution as ones using all seven (Figures 5A and B). Interestingly also, the distinction between the three strain groups at 31°C was greater in the later Days, whereas at 27°C it was greater in the earlier Days. Under these two conditions (i.e., later Days at 31°C and earlier Days at 27°C), the SS+ and WT groups were clearly distinct from one another in LD1/LD2 space, with the SS- group intermediate between the other two.

**Figure 5.**
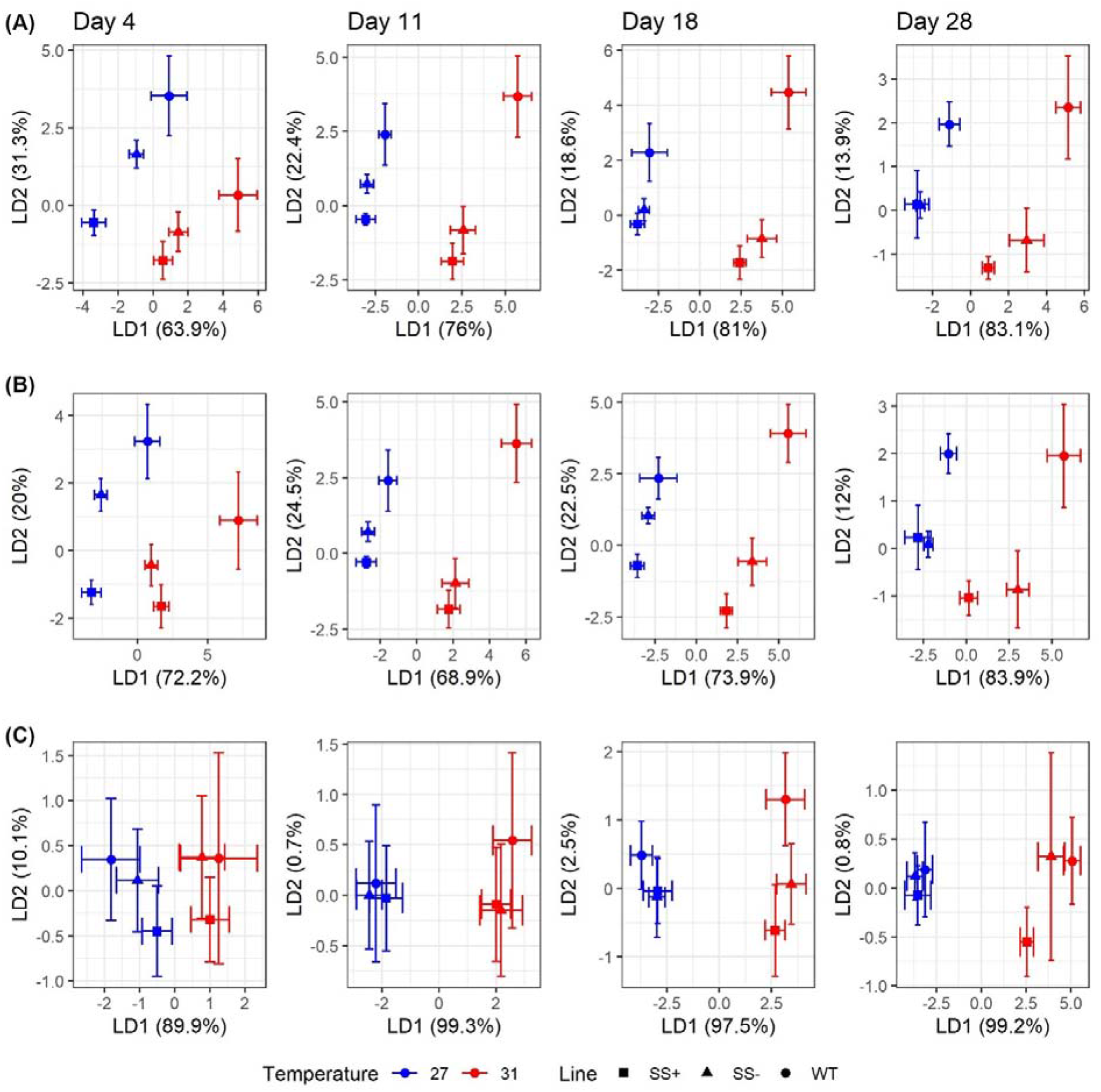
Plot of the two major linear discriminant functions for each strain type and temperature using various combinations of variables. **(A)** uses all seven retained variables (i.e., excluding ΦPSII and FSC), **(B)** just uses the five retained cytometry-based variables (i.e., ROS), ΔΨm, rGSH, Chl and SSC) and **(C)** just uses the two non-flow cytometry variables retained (i.e., CellDen and Fv/Fm). Mean values with their 95% confidence limits are given for the strain group (WT, SS+, SS-).

The LDAs only using the two non-flow cytometer variables retained, CellDen and Fv/Fm, showed broadly similar trends, including clearer discrimination between the strain groups in later Days at 31°C and earlier ones at 27°C, but overall, the resolution was somewhat less (Figure 5C).

The patterns in the LDAs thus showed that most of the variables contributed to some degree to the differentiation between strains. However, given the prominence of ROS, ΔΨm and rGSH in previous literature on heat tolerance, and the importance of ROS and ΔΨm revealed in our analyses, we also examined all the pairwise plots of those three variables alone. We found that most of the six pairwise plots for the two earlier Days at 27°C were able to distinguish the three types of strains, with no overlap between SS+ and WT and SS- again intermediate between them, albeit with some overlap with WT (Figure 6, S4).

**Figure 6.**
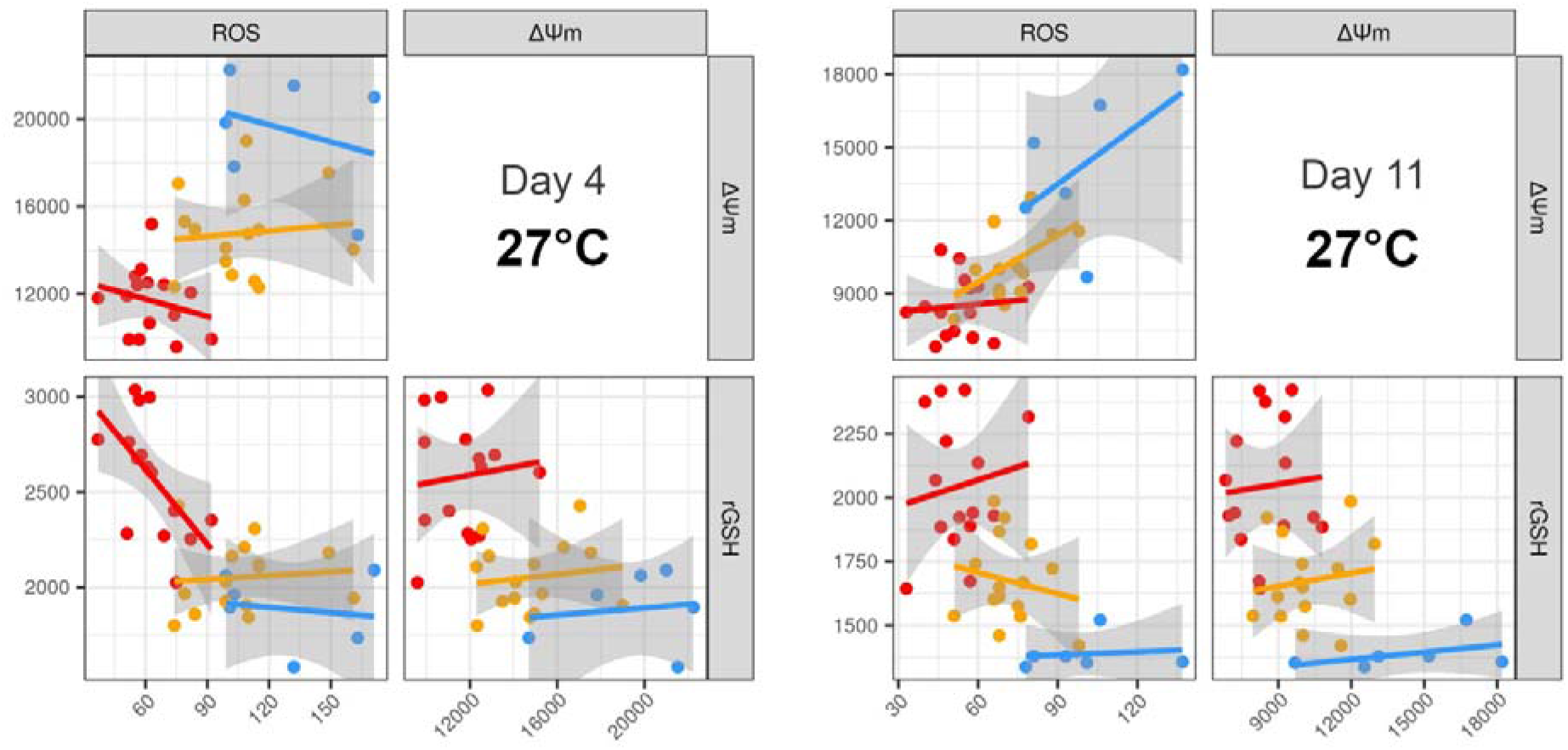
Pairwise plots of the three variables that most clearly separate the three strain types. The best resolution was achieved with a combination of ROS, ΔΨm and rGSH at ambient temperature at Days 4 and 11. WT (blue), SS- (yellow), SS+ (red). Each point on each plot represents a different replicate of the respective strains. Equivalent plots for Days 18 and 28 at each temperature are shown in Figure S4.

## DISCUSSION

### Differences between SS and WT strains

The six heat-evolved *Cladocopium* C1^acro^ SS strains we have studied had been exposed to 31°C for 9 years before this work. Our preliminary biochemical analyses after 6 years of exposure (Buerger *et al*., 2020) showed that they out-performed the WT in terms of their cell growth, photosynthetic performance and ROS secretion at 31°C. Those earlier results are supported and extended by the progress curves of the current experiment. The current results show that the adaptation of the SS strains to the elevated temperature is associated with a variety of physiological changes, some of them manifest at both ambient and elevated temperatures and some only in the latter.

Compared to WT, the SS strains had lower values for intracellular ROS and ΔΨm as well as higher rGSH at both temperatures (except for rGSH at 31°C) at the start of the experiment and throughout its the duration. This suggests some of the adaptation of the SS strains to the elevated temperature was based on constitutively expressed differences in genes affecting oxidant/antioxidant pathways. This idea is also borne out by our previous findings of higher constitutive gene expression of heat tolerance genes at ambient temperatures and lower secreted extracellular ROS in the SS strains at elevated temperatures (Buerger *et al*., 2020).

Genes affecting ΔΨm and rGSH levels are good candidates to be causally involved in the difference between the SS and WT strain groups. Evidence from various organisms under heat or other stresses has implicated lower mitochondrial membrane potentials and higher rGSH levels as mechanisms for decreasing ROS production and enhancing ROS scavenging, respectively (Owen and Butterfiel, 2010; Demine *et al*., 2019). One mechanism underpinning the lower ΔΨm could be a decoupling of the mitochondrial membrane potential and, as a result, reduced ATP production from the mitochondria (Demine *et al*., 2019), which could also lead to slower growth rates in the absence of compensating production elsewhere (Larosa and Remacle, 2018). We saw no evidence of this in the current study, but did see evidence for such a growth trade-off at ambient temperature in another experiment (Buerger *et al*., 2022). Also of note is that ΔΨm actually increased in later Days in all strains at both temperatures, but particularly in WT at 31°C. This may indicate some level of hyperpolarisation of the mitochondrial membranes, which can be an early sign of severe cellular stress (Weir *et al*., 2003) and would be consistent with the drop in cell densities observed, particularly in WT, in later Days at 31°C.

Other variables related to photosynthetic efficiency, cell density, size and morphology also differed between SS and WT strains. But these differences were generally more evident at 31°C than 27°C or at later rather than earlier sampling times. These temperature and time dependent changes might reflect genetic adaptations that are induced by the high temperature stress or consequential rather than primary, causal processes. In either event, the correlations among those variables and with ROS, ΔΨm and rGSH rarely explained more than half of their respective variances, so at least some of their differences reflect effects that were independent of each other and the oxidant/antioxidant differences.

Notably, comparative transcriptomics of three of the SS strains and the WT strain in holobionts with the coral *Acropora tenuis* at ambient temperatures had previously shown differences in the expression of genes involved in photosynthesis and measured photosynthetic yields between the two sorts of strains (Buerger *et al*., 2020). Notwithstanding the profound differences between *in vitro* and *in hospite* conditions, the two studies thus concur in finding differences between the SS and WT strains related to photosynthesis.

Our findings for FSC indicating that the adaptations of the SS strains to the elevated temperature included relatively smaller cell sizes than WT also corroborates some earlier literature. Thus exposure to elevated temperatures over several years has been found to result in smaller cell sizes for other microalgae, including *Emiliania huxleyi, Chrysochromulina rotalis* and *Prymnesium polylepis*, as well (Skau *et al*., 2017). However, other literature suggests it may not be a specific adaptation to heat stress; decreases in cell sizes have also been reported as adaptations to fluctuating high pCO_2_ stress that evolve over time in the green microalga *Ostreococcus* (Schaum *et al*., 2016). Evolution towards smaller cell sizes still may help the microalgae counteract a swelling effect of the stress on individual cells; such an effect has been reported for other Symbiodiniaceae strains exposed to 32°C (Fujise *et al*., 2018) and was evident in the progress curves for FSC values in all our strains, but particularly WT, over the 28 Day course of the experiment.

### Differences between SS+ and SS- strains

Importantly, our progress curves also showed differences in many of the variables between the SS+ and SS- strain groups, even though neither group had been exposed to elevated temperature in holobionts as opposed to *ex hospite* microalgae in culture since their establishment. In particular, ROS, CellDen, ΦPSII, Fv/Fm and SSC all showed clear differences between SS+ and both SS- and WT in the later time points at the higher temperature. Further, the performance of the SS- strains was in many cases intermediate between those of the SS+ and WT strains and the directions of the differences generally suggested that the SS+ strains experienced the least, WT the most, and SS- intermediate levels of stress at 31°C. These results implicate some of the same processes, particularly the oxidant/antioxidant and photosynthesis-related processes, operate to confer tolerance from microalgae to coral hosts in holobionts and *ex hospite* for microalgae in culture. Further, they suggest that more extreme changes from WT are needed to confer tolerance on the holobionts than from the heat-evolved microalgae.

Supporting the progress curves, the SS+ strains were also distinguished from the others in the temperature-dependent patterns of pairwise correlations among many of the variables. The most striking example involved cell densities, which showed little correlation with any other variable in SS+ at 31°C but were strongly correlated with most other variables in the other strain group-temperature combinations.

The key role of constituently expressed differences in oxidant/antioxidant-related variables in distinguishing SS+ from the other strain types is shown by the fact that they could be separated using the ROS, ΔΨm and rGSH data for 27°C alone. As noted, the differences in ROS measured here were in intracellular ROS, whereas in our preliminary work we did not find a difference between the heat selected strains in leaked, extracellular ROS (Buerger *et al*., 2020). Leaked extracellular ROS would impact holobiont tolerance through effects on the coral host whereas the intracellular ROS might do so through effects on the microalgal cell machinery. However, insomuch as it could represent the pool from which leaked ROS is obtained, the intracellular ROS therefore might also impact holobiont tolerance through effects on the host (Weis, 2008). We therefore suggest that cell integrity was maintained in all heat-evolved strains during elevated temperatures due to their relatively low level of secreted ROS. Compared to SS-, however, SS+ strains must have additional strategies to avoid intracellular ROS production to account for their increased thermal tolerance in holobionts.

The fact that the dye-based flow cytometry assays for ROS, ΔΨm and rGSH at 27°C could distinguish SS+ strains from the others we tested could also be significant from a practical point of view. If tests on other material previously characterised for holobiont conferring thermal tolerance produce the same pattern of differences, then these assays could prove a relatively rapid means of pre-screening large numbers of symbiont strains for candidates able to confer thermal tolerance on holobionts for use in reef restorations.

## Conclusions

We have found in our *in vitro* assessments that heat-evolved *Cladocopium* strains differ from an unselected control WT strain, derived from the same mother culture, in several aspects of oxidant/antioxidant levels, photosynthesis and cell size. The patterns of change suggest multiple genetic differences might have been involved, some constitutively expressed and some only evident at elevated temperatures. The more extreme phenotypes for some of these properties were also associated with higher temperature tolerance *in hospite*. In particular, expression of relatively low levels of intracellular ROS and ΔΨm and high levels of rGSH in *ex hospite* cultures at ambient temperature were found to be sufficient alone to distinguish the strains conferring tolerance *in hospite* from the others tested. If these differences prove generalisable, the flow cytometry based assay we developed here could prove a useful screening tool for symbiont strains of potential utility in the restoration of reefs bleached by high temperature stresses.

## Acknowledgements

The authors have no conflict of interest and no financial interest to declare. We thank Dr Melanie Carmody and Prof Barry Pogson from the Australian National University for providing the imaging pulse-amplitude modulation chlorophyll fluorometer for the photosynthetic measurements.

## Author Contributions

Conceptualization: MB, LC, PB, JO, MvO, OE.

Data curation: HY, PB, LC, MB.

Formal analysis: HY, PB, LC, JO, MB.

Funding acquisition: LC, OE, JO, MB, PB, MvO.

Investigation: LC, PB, MB.

Methodology: MB, LC, PB, JO.

Project administration: LC, JO, OE.

Resources: LC, PB, MB, MvO, OE.

Supervision: LC, JO.

Validation: LC, PB, MB, HY, JO.

Visualization: HY, PB, LC, JO, MB.

Writing – review & editing: PB, JO, LC, HY, MB, MvO, OE.

## Data availability statement

All raw data is provided along with the manuscript and as part of the supplementary materials.

## Ethics approval statement

Not applicable.

