## supplementary for "Flow cytometry-based biomarker assay for *in vitro* identification of heat tolerance conferring coral symbionts"

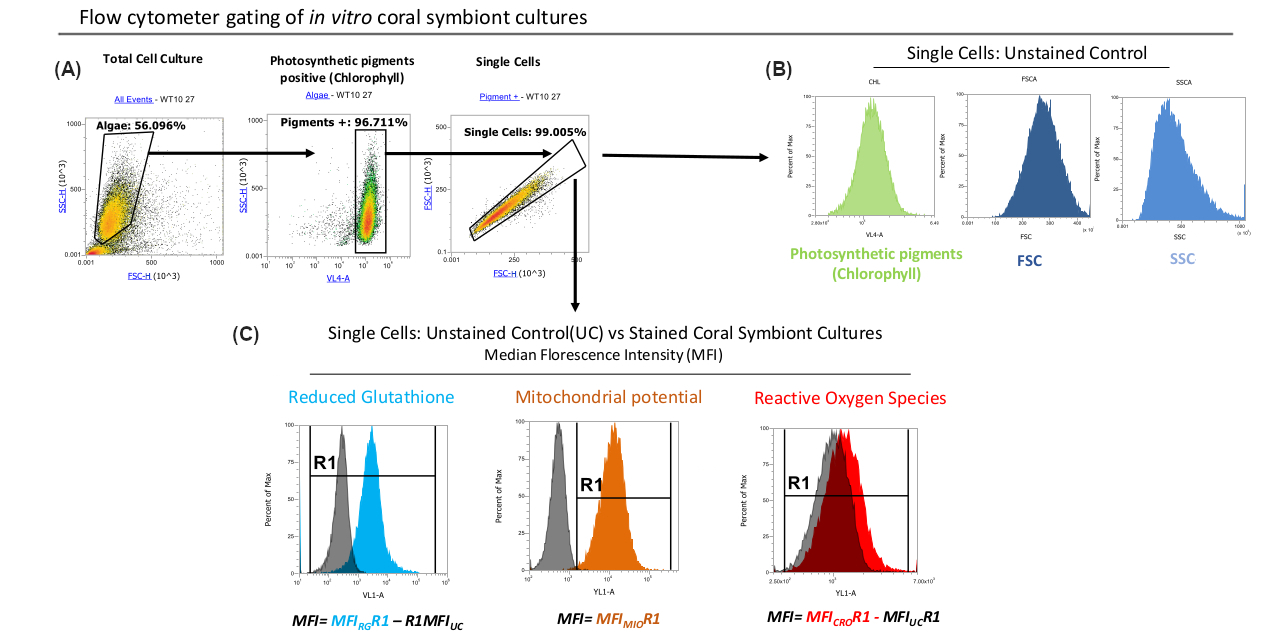

**Figure S1 Flow cytometry workflow. (A)** Microalgal cells were gated based on their characteristic FSC and SSC patterns. Only cells that were positive for photosynthetic pigments signal (likely chlorophyll) were selected for downstream analysis. **(B)** MFI values of Chlorophyll, FSC and SSC were measured in unstained controls (UCs). **(C)** Raw MFI values for rGHS and ROS were obtained in samples stained with Vita-Bright-48 and CellROX Orange reagents respectively and normalized by subtracting the autofluorescence signal from unstained controls in response to the excitation from the respective lasers. MFI values for CellROX Orange CMTMRos-stained cells used to measure mitochondrial potential (ΔΨm) were well separated from the autofluorescence signals of unstained controls and could therefore be analysed without normalisation against unstained controls.

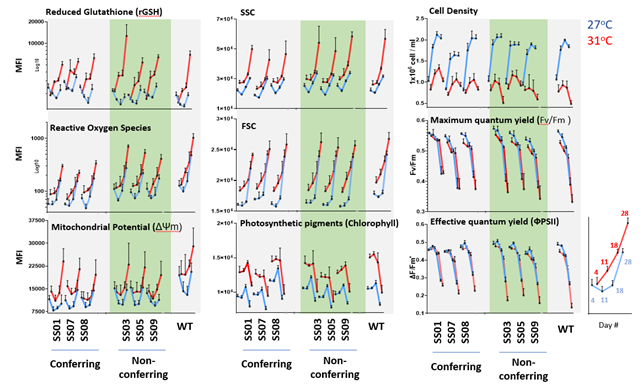

**Figure S2 Raw data projection.** Mean fluorescence intensities of microalgal cells over the course of the experiment. Dots represent consecutive sampling Days, red and blue lines indicate 31°C and 27°C assay cultures, respectively, and error bars show standard deviations. Numbers of replicate cultures were five each for the SS^+^ and SS^-^ strains and six for WT.

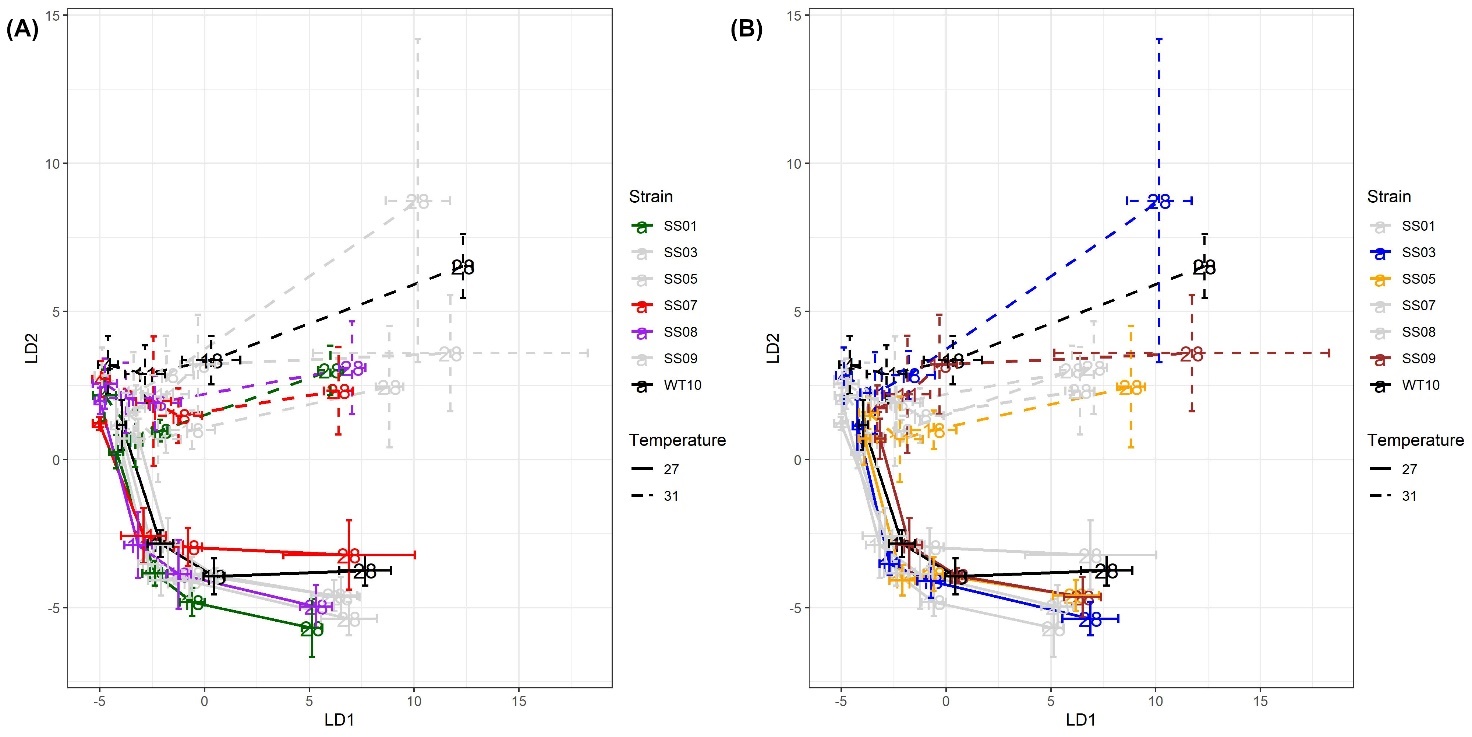

**Figure S3 LDA showing individual strain responses.** The same plot is shown in both panels, but different strains are highlighted. SS+ and SS- strains are highlighted in **(A)** and **(B),** respectively.

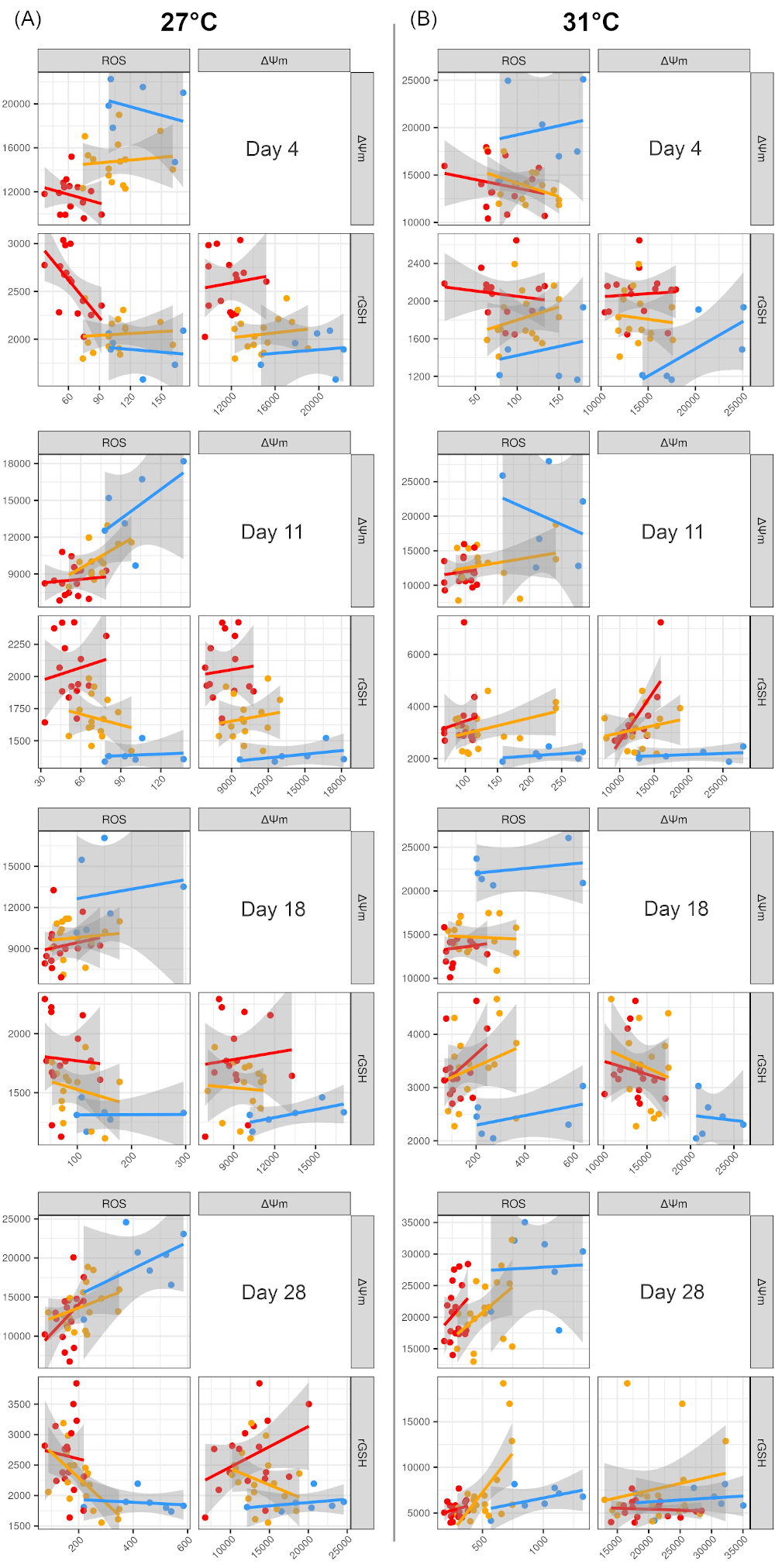

**Figure S4 Pairwise plots** **of ROS, rGSH and ΔΨm values for each Day and temperature.** WT, SS- and SS+ samples are shown in blue, yellow and red, respectively. Each point on each plot represents a different strain replicate.

**Table S1 Washing and staining conditions prior to flow cytometry.**

| Stain | Volume of cells [mL] normalized to 0.4x10^6^ cells / mL | Pre-stain washes * | Cell resuspension volume [mL] | Cell transfer volume [µL] | Final stain conc. [µM] | Staining time [min] | *Post-stain washes | Final cell re-suspension volume [µL] |
| --- | --- | --- | --- | --- | --- | --- | --- | --- |
| Vita-Bright-48 | 1 | 0 | - | - | 13.40 | 90 | 2 | 450 |
| Unstained Controls | 1.5 | 1 | 1.2 | 200 | - |  | 0 | 200 |
| CellROX  Orange |  | 1 |  | 500 | 3.00 | 60 | 0 | 500 |
| CMTMRos |  | 1 |  | 500 | 0.05 | 80 | 0 | 500 |

**Note: *** washes performed using 1mL RSS/IMK media at 2,600x g for 5 minutes.

**Table S2 Settings for reading stained algal cells on the flow cytometer**. * Unstained controls were also measured along with the respective dye and cytometer configurations to normalise the acquisitions of the dyed cells (Figure S1).

| Stain | Sample vol [µL] | Acquisition vol [µL] | Draw vol [µL] | Flow Rate  [µl/min] | STOP condition  [# of events on gate] | Stain excitation / emission maxima [nm] | Laser [nm] | Filter  [peak em / bandwidth] |
| --- | --- | --- | --- | --- | --- | --- | --- | --- |
| Unstained Controls | 200 | 90 | 150 | 500 | 20,000 cells | * | * | * |
| Vita-Bright-48 | 450 | 200 | 260 | 500 | 25,000 cells | 383/485 | 405 | 440/50 |
| CellROX Orange | 500 | 360 | 420 | 500 | 20,000 cells | 545/565 | 561 | 585/16 |
| CMTMRos | 500 | 370 | 430 | 500 | 15,000 cells | 554/576 | 561 | 585/16 |

**Table S3 Principal components.** Table shows the variance explained by the first seven PCs and the loadings of the various response variables on each PC. To simplify the model, ФPSII and FSC were removed from the analysis.

|  | **PC1 (54.2%)** | **PC2 (22.3%)** | **PC3 (8%)** | **PC4 (6.2%)** | **PC5 (3.9%)** | **PC6 (3.7%)** | **PC7 (1.7%)** |
| --- | --- | --- | --- | --- | --- | --- | --- |
| **CellDen** | -0.2628 | 0.5736 | -0.3110 | -0.4720 | 0.1426 | 0.5061 | 0.0769 |
| **Fv/Fm** | -0.4065 | -0.3521 | 0.0170 | 0.3350 | 0.4515 | 0.4502 | -0.4379 |
| **ROS** | 0.4417 | 0.1385 | -0.2522 | 0.0078 | 0.7958 | -0.2978 | -0.0108 |
| **ΔΨm** | 0.3983 | -0.1694 | -0.6360 | 0.3829 | -0.2432 | 0.4035 | 0.1988 |
| **rGSH** | 0.4242 | -0.0591 | 0.6242 | -0.0519 | 0.1727 | 0.5205 | 0.3514 |
| **Chl** | -0.0803 | -0.7031 | -0.2120 | -0.6204 | 0.1018 | -0.0193 | 0.2421 |
| **SSC** | 0.4749 | -0.0366 | 0.0170 | -0.3615 | -0.2079 | 0.1358 | -0.7620 |
| **Cumulative %** | 54.2% | 76.5% | 84.5% | 90.7% | 94.6% | 96.3% | 98% |

**Table S4 Linear discriminant analysis.** Table shows the variance explained by the respective LDs and the loadings of the various PCs on each LD.

|  | **LD1 (66.1%)** | **LD2 (23.3%)** | **LD3 (5.6%)** | **LD4 (3.4%)** | **LD5 (1.5%)** |
| --- | --- | --- | --- | --- | --- |
| **PC1 (54.2%)** | -1.8508 | -0.8895 | -0.0323 | 0.0206 | -0.0441 |
| **PC2 (22.3%)** | -2.0686 | 1.7082 | -0.4200 | 0.0970 | -0.0375 |
| **PC3 (8%)** | 0.7335 | -0.2339 | -0.1336 | 1.7139 | -0.5950 |
| **PC4 (6.2%)** | 1.5215 | -0.9796 | -2.3190 | -0.2152 | 0.0274 |
| **PC5 (3.9%)** | 0.3647 | 0.2045 | 0.0831 | -1.0344 | -2.0853 |
